# Wengen, a Tumour Necrosis Factor Receptor, regulates the Fibroblast Growth Factor pathway by an unconventional mechanism

**DOI:** 10.1101/2023.01.26.525651

**Authors:** Annalisa Letizia, Maria Lluisa Espinàs, Marta Llimargas

## Abstract

Unveiling the molecular mechanisms of receptor activation has led to much understanding of development as well as the identification of important drug targets. We use the *Drosophila* tracheal system to study the activity of two families of widely used and conserved receptors, the TNFRs and the RTK-FGFRs. Breathless, an FGFR, is known to respond to ligand by activating the differentiation program of the tracheal terminal cell. Here we show that Wengen, a TNFR, acts independently of both its canonical ligand and its downstream pathway genes to repress terminal cell differentiation. In contrast to Breathless, Wengen does not stably localise at the membrane and is instead internalised — a trafficking that seems essential for activity. We find that Wengen forms a complex with Breathless, and both colocalise in intracellular vesicles. Furthermore, Wengen regulates Breathless accumulation, likely regulating Breathless intracellular trafficking and degradation. We propose that, in the tracheal context, Wengen interacts with Breathless to regulate its activity in terminal cell differentiation. We suggest that such unconventional mechanism, involving binding by TNFRs to unrelated proteins, may be a general strategy of TNFRs activity.

## INTRODUCTION

Receptors receive information from the environment (e.g. as signalling molecules or mechanical forces) and transmit it to the cell to elicit changes. Their activity regulates many kinds of biological events during development and homeostasis, ranging from migration or cell differentiation to immunity or regulation of metabolism. Receptor activation needs to be exquisitely controlled to provide an outcome only when and where it is required. Thus, misregulation of receptor activity (excess or defect) frequently leads to malignant transformation, diseases or malformations.

Fibroblast Growth Factor Receptors (FGFRs), which belong to the Receptor Tyrosine Kinase (RTK) superfamily, are involved in diverse processes, ranging from organ morphogenesis to injury repair and regeneration. Consequently, FGFR malfunction leads to severe diseases, such as chronic kidney disease, dwarfism syndromes or obesity and it is also involved in cancer, especially in breast, lung, prostate and ovarian cancers ^1–3^. FGFRs are activated by Fibroblast Growth Factors (FGF) ligands. Ligand binding promotes receptor dimerisation and trans-phosphorylation, initiating the activation of downstream cascades, namely AKT, PLCγ, STAT and ERK-MAPK, by phosphorylation ^4^. Correlating with the importance of FGFRs in health and disease, their activity is finely regulated by a variety of mechanisms, including synthesis and secretion, stabilisation of FGF/FGFR, interactions with cofactors/adaptors, subcellular localisation, endocytosis and intracellular trafficking ^5^.

Tumor Necrosis Factor Receptors (TNFRs) also play key roles in development and homeostasis, and are particularly involved in the regulation of the immune system, inflammation and cell death. Missregulation of TNFR activity also leads to several serious pathologies such as autoinflammatory diseases and cancer ^6–8^. TNFRs are activated by Tumor Necrosis Factor (TNF) ligands, resulting in trimeric TNFR-TNF complexes. Through oligomerisation, the TNFR-TNF complexes recruit adaptor proteins like TRADD or TRAFs that initiate a cascade to regulate downstream signalling by JNK, NF-kB and Complex-II mediated apoptosis ^9,10^.

Because of the multifunctional nature of most receptor families in health and disease, it is urgent to understand the cross-talk between the different receptors, their signalling pathways, and their downstream outputs in “in vivo” conditions. *Drosophila* gives us many genetic tools in the approach to such complex problems. Here we use the embryonic tracheal system of *Drosophila* as a model to investigate the roles and interactions of two different types of receptors, the FGFR-Breathless (Btl), and the TNFR-Wengen (Wgn). The FGFR-Btl is central to tracheal development, regulating different steps including migration and cell differentiation (for reviews see ^11^,^12^). TNFR-Wgn was identified several years ago as a receptor for the unique TNF in *Drosophila* Eiger (Egr) ^13–16^, however its function, particularly in physiological conditions is unclear and controversial ^17^. Here we identify a physiological role for TNFR-Wgn in regulating the differentiation of tracheal cells. We show that TNFR-Wgn and FGFR-Btl are expressed in the same cells, regulate the same process, but do it in completely different ways. We find that TNFR-Wgn works in an unconventional manner to regulate the gradient of activity of FGFR-Btl, adding another layer of regulation of this critical receptor.

## RESULTS

### TNFR-Wgn is required to restrict tracheal terminal cell differentiation

We identified the TNFR-Wgn in the course of a genetic screen for new factors regulating tracheal development.

We used a null allele of *TNFR-wgn* (*TNFR-wgn^KO^* ^17^) to investigate *TNFR-wgn* tracheal requirements. The early steps of tracheal formation and branching were not affected (Fig S1A-F), however, we detected adventitious terminal branches throughout the whole tracheal tree (i.e. in dorsal branches (DBs), lateral trunk (LT), ganglionic branches (GBs) and visceral branches (VB)) (Fig 1A-D, S1A-F). To determine the origin of these terminal branches we stained the embryos with DSRF, a marker for terminal cell differentiation ^18,19^. We observed excess of DSRF positive cells that generated these adventitious terminal branches (Fig 1C-F). The *TNFR-wgn^KO^* phenotype was fully penetrant as all embryos displayed extra terminal cells. We focused on DBs, which in normal conditions contain 1 terminal cell at the tip (Fig 1E), to analyse the phenotype of *TNFR-wgn* depletion. We found around 90% of DBs containing more than one terminal cell, with a high proportion of them containing 3 or more (Fig S1G). In addition, we found many cases in which terminal cells also appeared in the stalk of the branch (Fig 1B,F, S1D,E,G). Quantification of terminal cells in other branches, like GBs, also indicated a significant increase with respect to the control (2,1% of GBs contained more than one terminal cell in the control, n=475 branches analysed, 57% of GBs contained more than one terminal cell in *TNFR-wgn^KO^* mutants, n=415 branches analysed).

**Fig 1:**
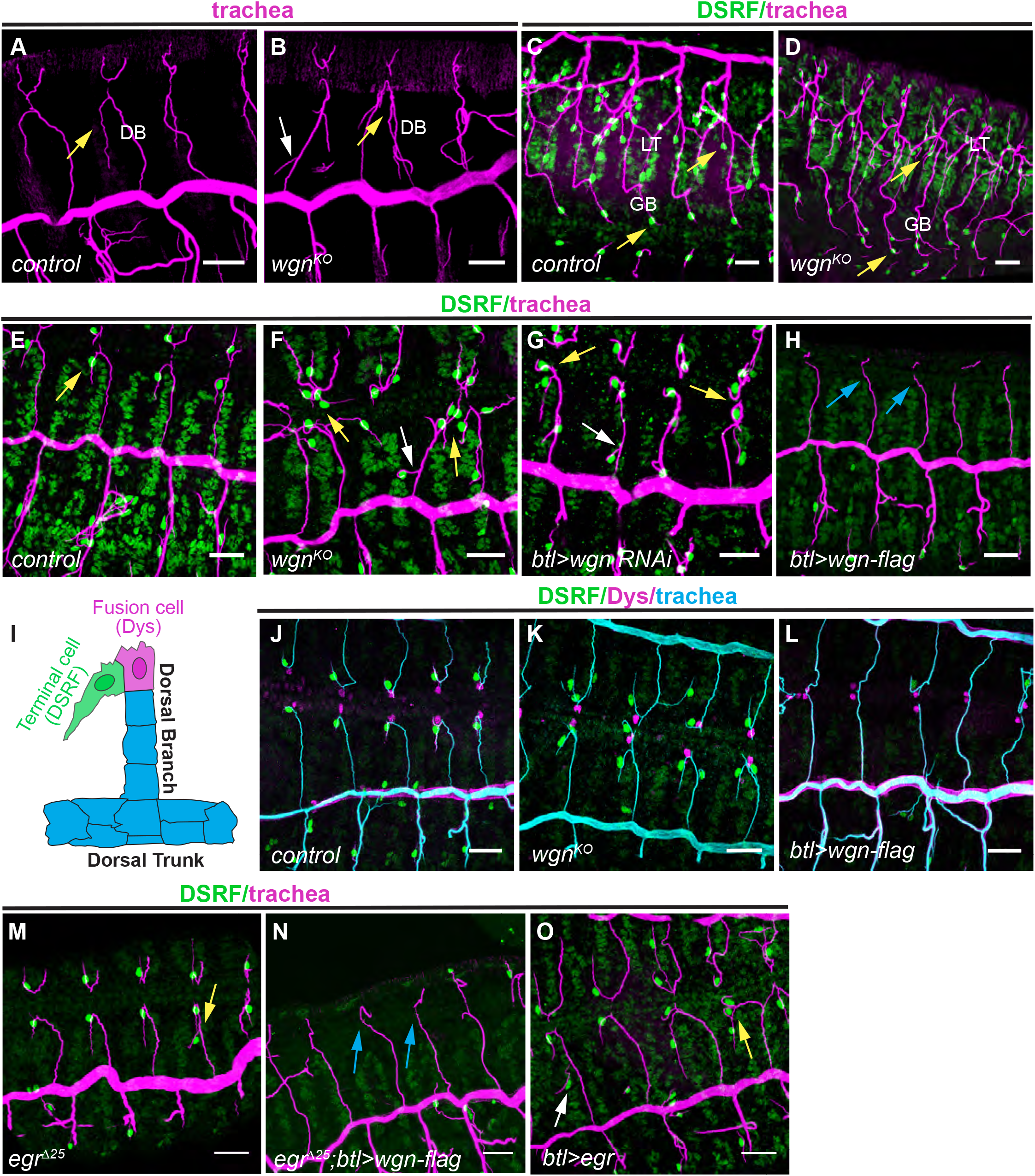
Functional requirements of *TNFR-wgn* and *TNFR-egr* during tracheal formation. (A-H) Dorso-lateral (A-B; E-H) or ventro-lateral (C,D) views of stage 15/16 embryos of the indicated genotypes stained with CBP to visualise the tracheal tubes (magenta) and with DSRF to visualise the terminal cells (green). Note the presence of one single terminal cell and terminal branch in the dorsal branches (DB) and in the ganglionic branches (GB) in wild type conditions (yellow arrows in A,C,E). In *TNFR-wgn* loss of function conditions more terminal branches and terminal cells arise from the tip of DBs (yellow arrows in B,F,G) or in the stalk (white arrows in B,F,G) and in GBs and Lateral trunk (LT) (yellow arrows in D). In conditions of *TNFR-wgn* overexpression terminal cells and terminal branches are not present (blue arrows in H). (I) Scheme showing the tip of a DB with one terminal cell expressing DSRF (forming a terminal branch) and one fusion cell expressing Dys engaged in branch fusion. (J-L) Dorsal views of stage 15 embryos of the indicated genotypes stained with CBP to visualise the tracheal tubes (cyan), with DSRF to visualise the terminal cells (green) and Dys to visualise the fusion cells (magenta). Note that fusion cells are normally specified in gain or loss of *TNFR-wgn* function. (M-O) Dorso-lateral views of stage 15/16 embryos of the indicated genotypes. *TNF-egr* mutants show a mild tracheal phenotype (yellow arrow in M) different from *TNFR-wgn* mutants (compare M to B,F). Overexpression of *TNFR-wgn* prevents terminal cell differentiation in the absence of *TNF-egr* (blue arrows in N). The overexpression of *TNF-egr* (O) leads to extra terminal cells (yellow arrows) also in the stalk (white arrow). Scale bar: 20 μm

Downregulation of *TNFR-wgn* in the tracheal system using RNAi reproduced the phenotype of null mutants (Fig 1G, S1G), indicating that TNFR-Wgn is required in the tracheal cells to regulate terminal cell differentiation.

The tracheal overexpression of a wild type form of *TNFR-wgn (TNFR-wgn-Flag*) ^20^ produced a highly penetrant phenotype that was the opposite to that shown by the loss of function: a loss of DSRF expressing cells (and terminal branches) throughout the tracheal system (Fig 1H, S1H).

*TNFR-wgn* manipulations specifically affected the differentiation of terminal cells, and neither the absence nor the overexpression of *TNFR-wgn* affected the differentiation of other tracheal cell types such as the fusion cells (Fig I-L).

Altogether these results show that *TNFR-wgn* activity is specifically required to limit the differentiation of the terminal cells that generate the terminal branches.

### TNFR-Wgn acts independently of its canonical signalling pathway during tracheal development

TNFR-Wgn was proposed to transduce the Egr signal through the JNK pathway ^21^. Thus, we investigated the contribution of this pathway to *TNFR-wgn* tracheal requirements (Fig 2A).

**Fig 2:**
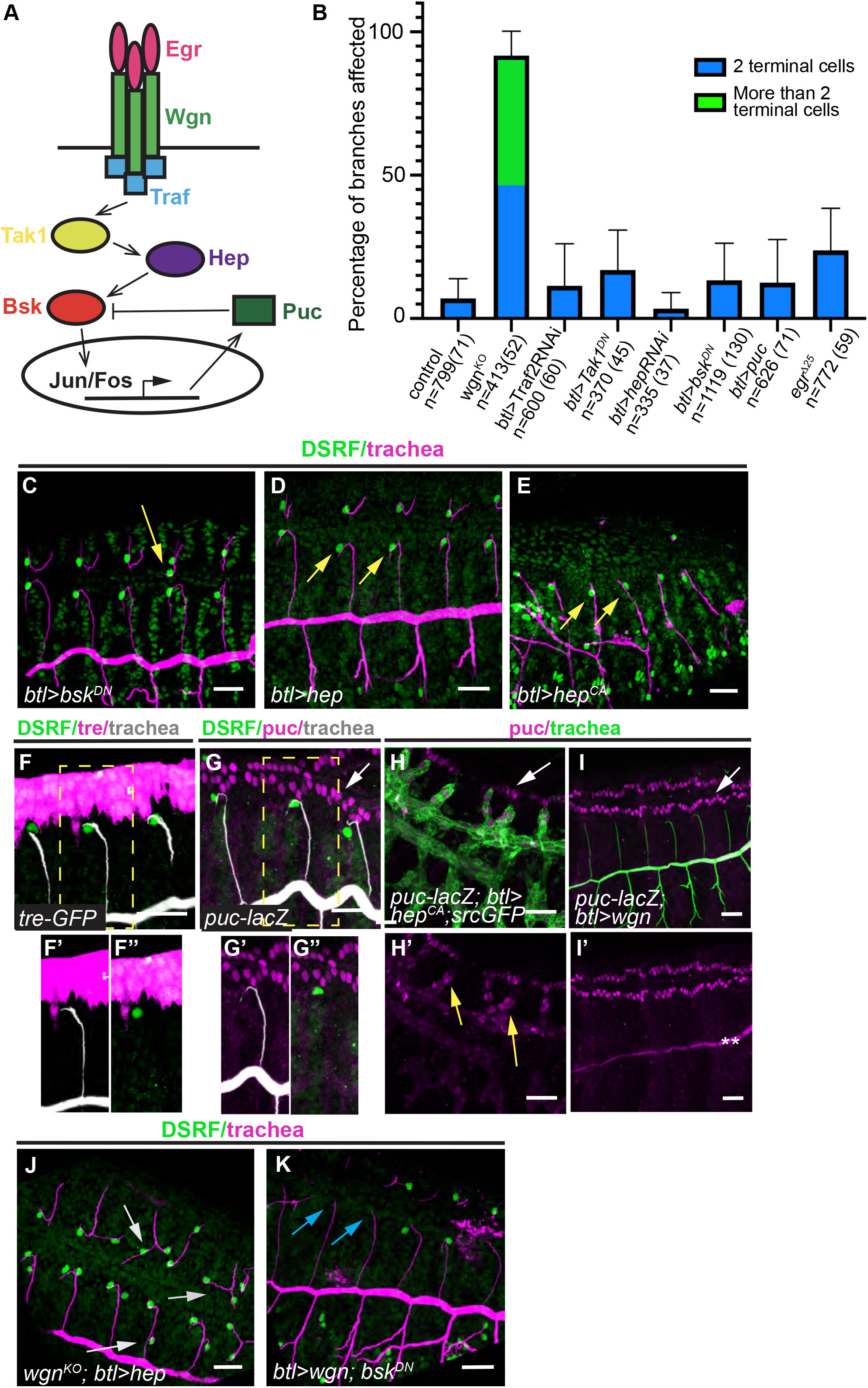
Analysis of JNK pathway role in terminal cell specification. (A) Scheme of the proposed Egr-Wgn-JNK signalling pathway (B) Quantification of the percentage of dorsal branches that show the indicated phenotypes in the different genotypes. Error bars indicate standard deviation (s. d.). n, number of DBs analysed, in brackets number of embryos analysed (C-E,J,K) Lateral views of stage 15 embryos of the indicated genotypes stained with CBP to visualise the tracheal tubes (magenta) and with DSRF to visualise the terminal cells (green). JNK pathway downregulation produce a mild tracheal effect (C). Note the presence of terminal cells in embryos with overactivated JNK (yellow arrows in D,E). Overexpression of *hep* cannot rescue the excess of terminal cells (grey arrowheads) produced by *TNFR-wgn* depletion (J). Downregulation of JNK cannot rescue the lack of terminal cells (blue arrowheads) produced by an excess of *TNFR-wgn* (K). (F,G) Lateral views of stage 14 embryos of the indicated genotypes stained with GFP or β-Galactosidase (magenta) to visualise the JNK activity reporters, with CBP to visualise the tracheal tubes (white) and with DSRF to visualise the terminal cells (green). Note that the reporters are not expressed in the trachea. White arrow in G points to *puc-lacZ* expression in the leading edge (H,I) Lateral views of stage 14 embryos of the indicated genotypes stained with β- Galactosidase (magenta) to visualise the *puc-lacZ* reporter, and in green to visualise the trachea. Note that *puc-lacZ* is activated in the trachea upon JNK overactivation (yellow arrows in H’) but not upon *TNFR-wgn* overexpression. Asterisks in J’ indicate staining background in the lumen of the trachea and white arrows in H,I point to *puc-lacZ* expression in the leading edge. Scale bar: 20 μm

Downregulation of the pathway using different tools (*bsk^DN^, Tak1^DN^, hepRNAi, Traf2RNAi*, or *UASpuc*) did not reproduce the *TNFR-wgn* loss of function phenotype. We never detected more than 2 terminal cells per DB, presence of terminal cells in the DB stalk, or a significant excess of terminal cells in GBs. We detected a low proportion of DBs with 1 extra-terminal cell (Fig 2B,C). Thus, the JNK downregulation phenotype did not correlate quantitatively and qualitatively with that of *TNFR-wgn* loss of function. In line with this result, overactivation of the pathway, using the overexpression of *hep* or a constitutively active form of *hep* (*hep^CA^*), did not prevent terminal cell specification (Fig 2D,E).

Different reporters that typically indicate JNK activity (*puc-lacZ* or *Tre-GFP*) were not detectably expressed in tracheal cells (Fig 2F,G), suggesting no role or a minor role of the pathway in the trachea. However, *puc-lacZ* tracheal expression was detected upon activation of the pathway with *hep^CA^* (Fig 2H). In contrast, upon overexpression of *TNFR-wgn, puc-lacZ* was not expressed in the tracheal cells (Fig 2I), indicating that *TNFR-wgn* does not detectably activate the pathway.

Finally, we found that the overexpression of *hep* cannot rescue the excess of terminal cells in *TNFR-wgn^KO^* mutants (Fig 2J), and that *bsk^DN^* did not revert the loss of terminal cells produced by *TNFR-wgn* overexpression (Fig 2K).

Altogether our results argue that *TNFR-wgn* does not act through the JNK pathway to regulate terminal cell differentiation.

### TNFR-Wgn acts independently of its canonical ligand during tracheal development

TNFR-Wgn was proposed to transduce the signal of the unique TNF in *Drosophila*, TNF-Egr ^14,15^. When we analysed null mutants for *TNF-egr* we could not detect DBs with more than 2 terminal cells, presence of terminal cells in the stalk, or a significant excess of terminal cells in GBs as in *TNFR-wgn* mutants. We detected a low penetrant phenotype of 1 extra-terminal cell in DBs (Fig 1M), similar to the effects of JNK pathway downregulation (Fig 2B). These quantitative and qualitative phenotypic differences strongly suggested that TNFR-Wgn regulates terminal cell differentiation independently of its ligand TNF-Egr, although we cannot completely discard a minor contribution. In agreement with this hypothesis, we found that the overexpression of *TNFR-wgn* was still able to prevent terminal cell specification in the absence of *TNF- egr* (Fig 1N).

We also investigated the effect of *TNF-egr* overexpression in the trachea. Instead of producing a phenotype comparable to that of *TNFR-wgn* overexpression (i.e. absence of terminal cells), it produced an effect comparable to the *TNFR-wgn* loss of function (i.e. significant excess of terminal cells, also in the DB stalk) (Fig 1O). This result fits in a model in which TNFR-Wgn acts independently of TNF-Egr in the trachea. In this scenario, the presence of TNF-Egr in the trachea would bind and sequester TNFR-Wgn, preventing it from performing its independent physiological activity (see below).

### TNFR-Wgn accumulates in intracellular vesicles in the tracheal cells

*TNFR-wgn* is expressed in different tissues, including the tracheal system (BDGP). We analysed the accumulation and localisation of TNFR-Wgn protein in tracheal cells. In contrast to our expectations for a membrane receptor, we could not detect TNFR-Wgn in the membrane of tracheal cells. Instead, we detected Wgn in intracellular punctae (Fig 3A,B), as previously described in imaginal discs ^22^. We reasoned that maybe the endogenous levels of TNFR-Wgn were not high enough to be detected at the membrane by the antibody. For this reason, we overexpressed TNFR-Wgn in the tracheal cells. We detected increased levels of TNFR-Wgn in these conditions, but again the protein localised moslty in intracellular punctae (Fig 3C). However, we also observed, on occasions, some accumulation of TNFR-Wgn in the apical membrane upon overexpression (Fig 3D). This result suggested that TNFR-Wgn has some ability to localise to the membrane, at least when we saturate the system.

**Fig 3:**
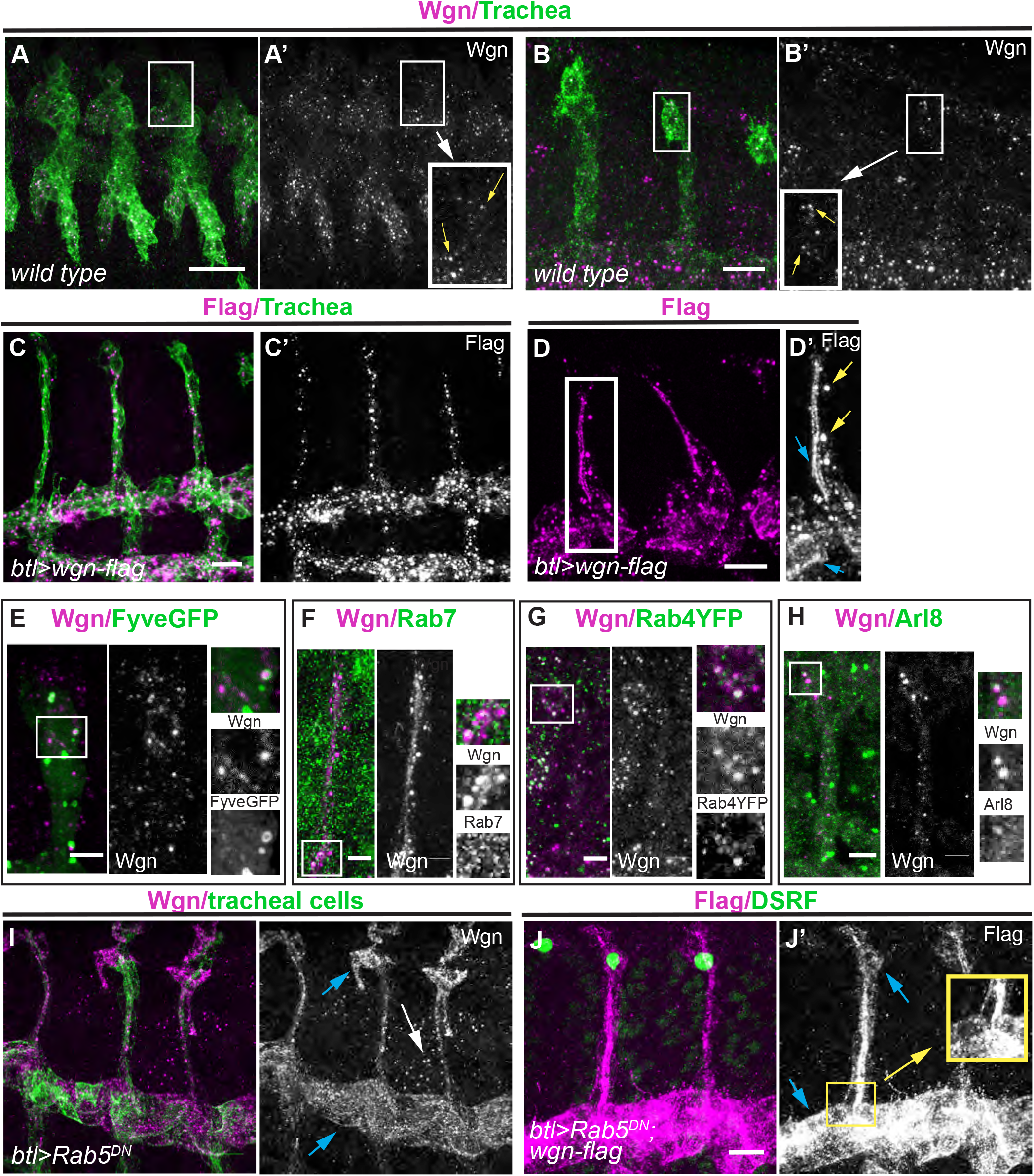
Pattern of TNFR-Wgn protein in the tracheal system. (A-J) All images show dorso-lateral views of stage 14 embryos focused to visualise the DBs, except A, which shows a general view of a stage 13 embryo. TNFR-Wgn (in magenta) is visualised using a Wgn antibody or a Flag antibody that recognises overexpressed *TNFR-wgn-Flag*. TNFR-Wgn accumulates in the tracheal cells in wild type conditions in intracellular vesicles (yellow arrows in insets in A,B). TNFR-Wgn overexpression shows a similar pattern (C), although, besides the accumulation in vesicles (yellow arrows in D’), accumulation in the apical region can be detected (blue arrows in D’). (E-H) Colocalisation analysis indicates that TNFR-Wgn colocalises with late endosomal markers (Fyve-GFP and Rab7), with the fast recycling marker Rab4 and with lysosomal markers (Arl8). (I-J) When internalisation is compromised in tracheal cells, endogenous or overexpressed TNFR-Wgn localises to the membrane (blue arrows in I,J), although TNFR-Wgn still accumulates in punctae in other tissues (white arrow in I). In these conditions, overexpression of TNFR-Wgn cannot prevent terminal cell differentiation (note presence of terminal cells in J) Scale bar: A 20 μm; B,C,D,I 10 μm; E-H 5 μm

To identify the nature of TNFR-Wgn punctae, we co-stained with different intracellular trafficking markers. We found colocalisation with markers of late endosomes and multivesicular bodies (i.e. Rab7 and Hrs, Fig 3E,F), with lysosomal markers (Arl8, Fig 3H) and with the fast recycling pathway component Rab4 (Fig 3G). Thus, the results indicated that TNFR-Wgn is internalised, traffics through the endocytic pathway and it is then degraded or recycled back to the membrane. Because we found this endocytic trafficking of TNFR-Wgn but could not detect TNFR-Wgn stabilised at the membrane, we speculated that TNFR-Wgn is constitutively internalised. To test this hypothesis, we compromised endocytic uptake by downregulating Rab5 activity. In these conditions, we now found a clear accumulation of TNFR-Wgn (endogenous and overexpressed) at the apical, basal, and lateral membrane (Fig 3I,J). Strikingly, we also found that when internalisation is compromised, TNFR-Wgn can no longer prevent terminal cell differentiation (Fig 3J), arguing that TNFR-Wgn must be internalised to exert its activity.

Altogether these results show that TNFR-Wgn is a highly dynamic protein that reaches the membrane but is constantly internalised, preventing it to stably localise at the membrane.

### TNFR-Wgn regulates the activity of the FGFR-Btl pathway

We asked how TNFR-Wgn regulates terminal cell differentiation.

Terminal cell differentiation was previously shown to be activated by FGF-Branchless (Bnl)/FGFR-Btl ^23,24^, which transduces the signal through the ERK-MAPK cascade ^25^ (Fig S2A). We asked whether TNFR-Wgn was regulating this signalling pathway and we found it did. In wild type conditions FGFR-Btl activation leads to phosphorylation of ERK that enters the nucleus and activates the terminal cell program in the tip cell (Fig 4A). We found that in *TNFR-wgn* mutant conditions many more cells accumulated dpERK in the nucleus, correlating with the excess of DSRF cells (Fig 4B). In contrast, in *TNFR-wgn* overexpression conditions, no dpERK accumulated in the nucleus of tip cells, correlating with absence of DSRF expressing cells (Fig 4C). These results indicated that TNFR-Wgn restricts terminal cell differentiation by downregulating the ERK-MAPK cascade activated by FGF-Bnl/FGFR-Btl.

**Fig 4:**
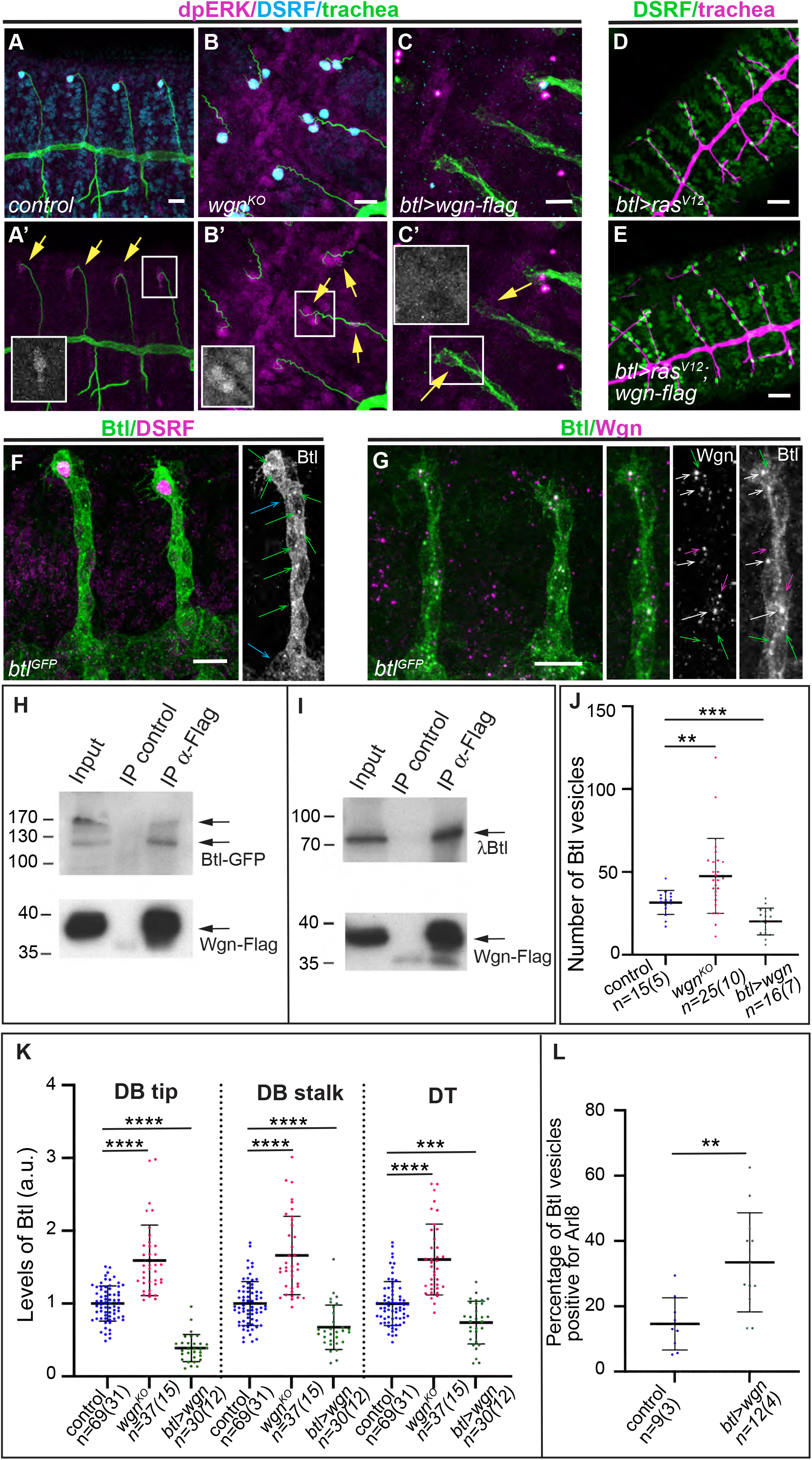
TNFR-Wgn forms a complex with FGFR-Btl and regulates its activity. (A-C) Dorso-lateral views of stage 14/15 embryos of the indicated genotypes stained with dpERK (magenta), DSRF (cyan) and a tracheal marker (green). dpERK accumulates in the nuclei of tip cells and activates DSRF. In *TNFR-wgn* mutants more cells accumulate nuclear dpERK, while in *TNFR-wgn* overexpression no dpERK accumulates in tip tracheal cells. (D-E) Dorso-lateral views of stage 14/15 embryos of the indicated genotypes stained with DSRF (green) and CBP to visualise the trachea (magenta). (F,G) Dorso-lateral views of a stage 14 embryos showing two dorsal branches stained to detect FGFR-Btl (green) and DSRF or TNFR-Wgn (magenta). Note the accumulation of FGFR-Btl at the basal membrane (blue arrows in F) and also in intracellular vesicles (green arrows in F). Many of the FGFR-Btl containing vesicles contain also TNFR-Wgn (white arrows in G), but TNFR-Wgn and FGFR-Btl single vesicles are also detected (magenta and green arrows respectively). (H,I) Co-immunoprecipitation experiments. Western blot using αBtl (upper panels) and αFlag (lower panels) of salivary gland extracts from third-instar larvae that express either *TNFR-wgn-Flag* and *FGFR-btl-GFP* (H) or *TNFR-wgn-Flag* and *FGFR-λbtl* (I). Note that αBtl recognizes two specific bands. (J-L) Scatter plots quantifying the number of FGFR-Btl intracellular vesicles (J), the levels of FGFR-Btl accumulation at the tip or stalk of the DBs or in the DT (K), or the proportion of FGFR-Btl intracellular vesicles that are positive for Arl8. Genotypes are indicated. n, number of DBs or DTs analysed, in brackets number of embryos analysed. Bars show mean ± SD. ****p < 0.0001; 0.0001<***p<0.001, 0.001 < **p<0.01, unpaired t test with Welch’s correction. Scale bar: A-C,F,G 10 μm; D,E 20 μm

Constitutive activation of Ras leads to supernumerary terminal cells (Fig 4D). This phenotype was not reverted when simultaneously overexpressing TNFR-Wgn (Fig 4E), suggesting that TNFR-Wgn acts upstream or in parallel to Ras.

### TNFR-Wgn forms a complex with FGFR-Btl receptor

As TNFR-Wgn seemed to act upstream of Ras, we considered the possibility that it regulates the FGFR-Btl receptor. To investigate this possibility, we used tagged alleles of *FGFR-btl* (*FGFR-btl^GFP^* and *FGFR-btl^endoRFP^*). *FGFR-btl-tagged* forms reproduced the pattern of expression of the gene (with higher levels at the tip during the specification of tip cells, ^26^, Fig S2B,C) and showed that the protein accumulated at the cell membrane, with a clear presence at the basal membrane (Fig 4F, S2B,C) from where the ligand FGF-Bnl is received ^24,27^. In addition, a detailed subcellular analysis detected FGFR-Btl in intracellular vesicles (Fig 4F). These vesicles likely reflect the normal intracellular trafficking and recycling of FGFR-Btl to ensure its proper localisation and activity ^5,28,29^. Co-staining with TNFR-Wgn indicated that many of these FGFR-Btl vesicles also contained TNFR-Wgn (Fig 4G, S2H). The presence of the two receptors in the same vesicles could just indicate that they traffic together, but it could also indicate a more direct interaction.

To test a possible interaction between TNFR-Wgn and FGFR-Btl we performed co-IP experiments. We expressed TNFR-Wgn and FGFR-Btl in salivary glands (which normally do not express these genes) and we found that the two proteins also colocalised in intracellular vesicles (Fig S2D). TNFR-Wgn co-immunoprecipitated full length FGFR-Btl as well as a constitutively active form of FGFR-Btl in which the extracellular domain has been replaced with the dimerization domain of the bacteriophage λ, λBtl (Fig 4H,I). These results show that TNFR-Wgn and FGFR-Btl form a complex and that the transmembrane and/or the intracellular domains of FGFR-Btl are sufficient for this interaction.

In a previous section we have hypothesised that the overexpression of TNF-Egr produced a *TNFR-wgn* loss of function phenotype because TNF-Egr interferes with a TNF-Egr-independent activity of TNFR-Wgn. In agreement with this hypothesis, we found that the proportion of common FGFR-Btl/TNFR-Wgn vesicles decreased when we overexpressed TNF-Egr (Fig S2H).

### TNFR-Wgn regulates FGFR-Btl accumulation

As we found that TNFR-Wgn forms a complex with FGFR-Btl and regulates its downstream activity, we asked whether TNFR-Wgn regulates FGFR-Btl at the cellular level. We found that this was the case. We first measured the levels of FGFR-Btl in tracheal cells. We observed a clear increase of FGFR-Btl levels in *TNFR-wgn* mutants, which was strongly detected at the basal membrane (Fig 4K, S2E,F). In contrast, we found a clear decrease of FGFR-Btl upon *TNFR-wgn* overexpression, which was very conspicuous particularly at the membrane of the tips (Fig 4K, S2E,G). We then quantified the presence of FGFR-Btl intracellular vesicles. We detected a significant increase of FGFR-Btl vesicles in *TNFR-wgn* mutants and a decrease in *TNFR-wgn* overexpression conditions (Fig 4J).

Thus, TNFR-Wgn regulates the general levels of FGFR-Btl accumulation and the presence of FGFR-Btl vesicles. Our results could indicate a role for TNFR-Wgn in promoting FGFR-Btl degradation. In this scenario, lack of TNFR-Wgn activity would lead to decreased FGFR-Btl degradation resulting in more vesicles and higher FGFR-Btl levels. In contrast, TNFR-Wgn overexpression would promote degradation resulting in less vesicles and lower levels. To test this possibility, we analysed the colocalisation of FGFR-Btl vesicles with Arl8 as a marker for lysosomes ^30^ in conditions of TNFR-Wgn overexpression. While there was a certain variability, we detected a significant increase of FGFR-Btl vesicles positive for Arl8 (Fig 4L), strongly suggesting that TNFR-Wgn regulates FGFR-Btl trafficking and promotes its degradation.

### TNFR-Wgn modulates the effects of active FGFR-Btl

To investigate whether TNFR-Wgn regulates terminal cell differentiation by regulating FGFR-Btl levels, we asked whether the overexpression of *FGFR-btl* could bypass the effect of *TNFR-wgn* overexpression. We co-overexpressed both *TNFR-wgn* and *FGFR-btl-GFP* (which rescues the lack of FGFR-Btl activity ^31^ and undergoes normal trafficking ^28^). We found a phenotype of lack of terminal cells (Fig S3A), indicating that the overexpression of *FGFR-btl* cannot rescue the defects produced by excess of *TNFR-wgn*. In parallel, we also found that the overexpression of *FGFR-btl-GFP* alone was not able to produce an excess of terminal cells (Fig S3B). Collectively, these results suggested that terminal cell differentiation does not depend on the total levels of FGFR-Btl, but may rather depend on the levels of activated FGFR-Btl. In agreement with this, we observed that the overexpression of *FGF-bnl* (Fig S3C) or activated *FGFR-λbtl*, (Fig 5A) led to extra terminal cells. We then asked whether the overexpression of *FGFR-λbtl* could bypass the effect of *TNFR-wgn* overexpression, and we found it did. The co-expression of *FGFR-λbtl* and *TNFR-wgn* produced a rescue of the lack of terminal cells (Fig 5B,C).

**Fig 5:**
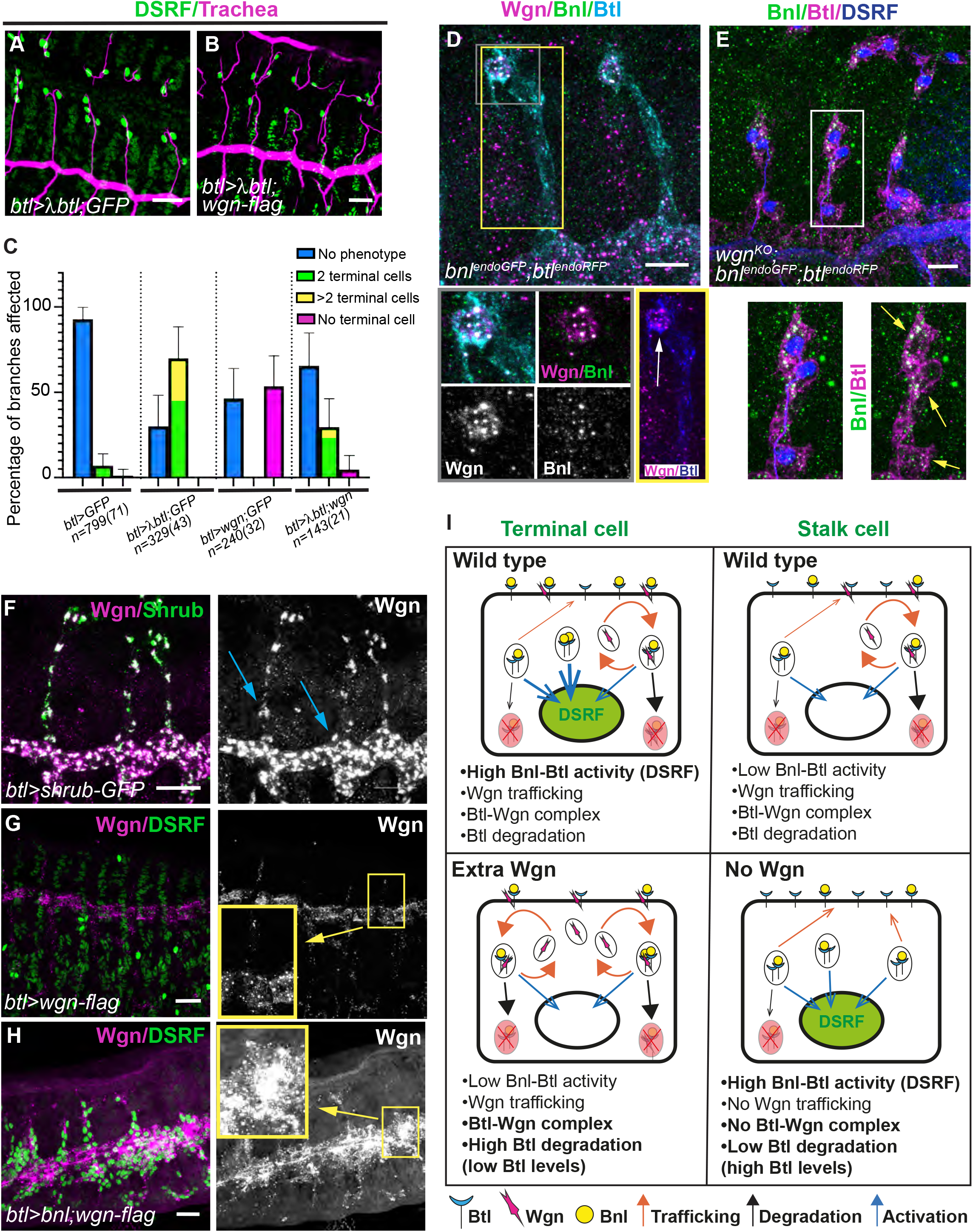
Interactions TNFR-Wgn/ FGFR-Btl. (A-B) Dorso-lateral views of stage 15 embryos of the indicated genotypes stained with DSRF (green) and CBP as a tracheal marker (magenta). (C) Quantification of the percentage of dorsal branches that show the indicated phenotypes in the different genotypes. Note the phenotype of extra terminal cells when FGFR-Btl is activated and the lack of terminal cells when TNFR-Wgn is overexpressed. The combination of the two conditions results in a rescue of each phenotype. Error bars indicate standard deviation (s. d.). n, number of DBs analysed, in brackets number of embryos analysed. (D-H) Dorso-lateral views of a stage 14 embryos of the indicated genotypes. (D) FGF-Bnl accumulates in intracellular vesicles that also contain TNFR-Wgn and FGFR-Btl particularly in the terminal cells (grey square). More TNFR-Wgn vesicles are detected at the tip of DBs (white arrow) compared to the base (yellow square). (E) In *TNFR-wgn* mutants, FGF-Bnl is detected in more cells of the DBs (yellow arrows). (F) When the endocytic maturation is compromised, TNFR-Wgn is found also at high levels in the stalk cells at the base (blue arrows). (G,H) The overexpression of FGF-Bnl leads to an increased accumulation of TNFR-Wgn protein. (I) Model. TNFR-Wgn protein is constantly trafficking and can form a complex with FGFR-Btl receptor. Through this interaction, TNFR-Wgn promotes FGFR-Btl degradation. In the wild type, the terminal cell receives huge amounts of FGF-Bnl ligand (due to the proximity to its source), activating FGFR-Btl, which leads to activation of the ERK pathway and to transcriptional DSRF activation. This activation can bypass the negative effect of TNFR-Wgn. The stalk cell receives lower levels of FGF-Bnl ligand, leading to a weaker activation of the pathway. This, combined with the negative effect of TNFR-Wgn, prevents DSRF activation. When TNFR-Wgn is overexpressed, TNFR-Wgn promotes FGFR-Btl degradation, preventing DSRF activation in spite of the presence of high levels of FGF-Bnl ligand. This also leads to lower levels of FGFR-Btl. When TNFR-Wgn activity is lost, FGFR-Btl degradation decreases, and FGFR-Btl levels increase. Under these conditions, a weak activation of the FGFR-Btl in stalk cells can lead to DSRF activation. Scale bar: A,B,F-H 20 μm; D,E 10 μm

To further explore the role of TNFR-Wgn in regulating active FGFR-Btl, we analysed FGF-Bnl distribution in tracheal cells using an endogenously tagged-*bnl* allele (*bnl^endoGFP^*, ^32^). In wild type conditions we observed the pattern of FGF-Bnl punctae close to the tracheal cells ^27^. In addition, we also detected FGF-Bnl accumulated in large and conspicuous intracellular vesicles containing also FGFR-Btl and TNFR-Wgn; these were mainly in the terminal cell (Fig 5C). This intracellular FGF-Bnl signal may correspond to the internalisation of active ligand-receptor complexes that signal through ERK to trigger the terminal cell program. In contrast to the wild type, we found the presence of FGF-Bnl intracellular vesicles in several cells of dorsal branches in *TNFR-Wgn* mutants which activate DSRF (Fig 5D). The results suggest that TNFR-Wgn restricts the accumulation/maintenance of FGF-Bnl/FGFR-Btl complexes, and likely FGFR-Btl activation, to more proximal regions in the branch.

### FGFR activity regulates TNFR-Wgn intracellular trafficking

We noticed that, during terminal cell differentiation, intracellular vesicles of TNFR-Wgn were more abundant in the terminal cells compared to the proximal part of the DB (Fig 3B, 5D). This result could indicate a faster or higher degradation of TNFR-Wgn at proximal regions and/or increased accumulation (transcriptional or posttranscriptional) at the tips. To investigate this further, we blocked endocytic maturation of vesicles using *shrub-GFP* ^33,34^. We found the presence of large vesicles containing TNFR-Wgn at both tip and proximal regions (Fig 5F), suggesting that in normal conditions TNFR-Wgn is differentially processed throughout the dorsal branch, and likely degraded faster at the proximal region, probably contributing to the TNFR-Wgn pattern.

Is TNFR-Wgn protected from degradation at the tips? As we had found that at the tips TNFR-Wgn intracellular vesicles contained FGF-Bnl, we asked whether FGFR-Btl activation could affect TNFR-Wgn trafficking. We found that in conditions of FGF-Bnl overexpression, overexpressed TNFR-Wgn was significantly increased in intracellular vesicles compared to control (Fig 5G,H). These results point to a regulatory feed-back loop in which FGF-Bnl presence (or activated FGFR-Btl receptor) stabilises TNFR-Wgn protein, while TNFR-Wgn regulates FGFR-Btl activity.

## DISCUSSION

We evidence a model in which two different receptors, each acting by a different mechanism, regulate one physiological event. It was previously known that FGFR-Btl, by FGF-Bnl ligand activation, regulates the differentiation of tracheal terminal cells. Here we find that the TNFR-Wgn also regulates this process and propose that it does so by directly regulating the activity of FGFR-Btl. Our biochemical data shows that TNFR-Wgn and FGFR-Btl form a complex. In addition, our cellular analysis shows that while TNFR-Wgn can localise at the membrane, in normal conditions it is constantly and rapidly internalised into intracellular vesicles. Furthermore, we demonstrate that TNFR-Wgn regulates FGFR-Btl accumulation. Therefore, our results strongly suggest that through its constant trafficking and its ability to interact with FGFR-Btl, TNFR-Wgn regulates the activity of FGFR-Btl in terminal cell differentiation.

Our biochemical data indicates that TNFR-Wgn forms a complex with activated and wild type FGFR-Btl receptors. However, our genetic experiments revealed an interaction between TNFR-Wgn and the activated FGFR-λBtl receptor, but not with the wild type FGFR-Btl: our results indicate that forced FGFR-Btl expression cannot overcome the repressor effect of TNFR-Wgn, suggesting that TNFR-Wgn restricts terminal cell differentiation by regulating FGFR-Btl activity rather than the absolute levels of FGFR-Btl. Moreover, the overexpression of wild type FGFR-Btl does not lead to extra terminal cell differentiation, strongly suggesting that the levels of the receptor are not limiting. In this respect, the haploinsufficient nature of the FGF-Bnl loss of function ^24^ suggests than the limiting factor for FGFR-Btl activation is the ligand availability. We propose that TNFR-Wgn interacts with activated and non-activated FGFR-Btl receptors, but this interaction only has phenotypic consequences for terminal cell differentiation in the case of the activated receptor. Several reports have shown that, although FGFRs can dimerise and internalise in the absence of ligand, FGFRs activation stimulates its endocytosis (reviewed in ^2^). Thus, it is possible that the activated FGFR-Btl (either *FGFR-λbtl* or when activated by *FGF-Bnl*) is involved in a more dynamic trafficking, and this could facilitate interactions with TNFR-Wgn, which also undergoes a dynamic trafficking.

How does TNFR-Wgn regulate FGFR-Btl activity? It is known that the strength, the duration and also the subcellular localisation of activated FGFRs can determine the cellular outcome (reviewed in ^35^). For instance, internalisation of activated FGFR1 does not attenuate the signal but instead promotes stronger signalling through the ERK pathway, while AKT activation is independent of FGFR internalisation ^36,37^. In our system, we propose a model (Fig 5I) in which FGFR-Btl is activated by the presence of FGF-Bnl at the tips, which stimulates its internalisation and signalling from endosomes through the ERK pathway to regulate the differentiation of terminal cells. As we find that TNFR-Wgn forms a complex with FGFR-Btl and promotes its degradation, TNFR-Wgn could be regulating the intensity or duration of internalised FGF-Bnl/FGRF-Btl complexes that traffic in endosomes, modulating ERK activation. Alternatively, TNFR-Wgn could be promoting FGFR-Btl internalisation and its degradation modulating the availability of receptor to respond to ligand binding. Thus, TNF-Wgn, by fine-tuning FGFR-Btl activity, would restrict the differentiation of terminal cells in tracheal branches. Finding new regulators of FGFR internalisation, trafficking and activity is critical for the development of new FGFR-directed therapies for disease and cancer treatments ^2,3,5^.

TNFR-Wgn was identified as the receptor for a unique TNF in Drosophila, TNF-Egr ^13–16^. However, this finding was subsequently questioned: it was shown that TNFR-Wgn can act independently of TNF-Egr in photoreceptor axon pathfinding ^20^. In addition, complicating the issue, a second TNFR was identified, named TNFR-Grindelwald (Grnd), and it was proposed that this is the receptor that transduces TNF-Egr functions ^17^. In a further development it has now been demonstrated that TNF-Egr can bind both TNFRs, TNFR-Grnd and TNFR-Wgn, but with very different affinities. TNFR-Grnd binds TNF-Egr with a much higher affinity than TNFR-Wgn, suggesting they have different cellular functions ^22^. Thus, a role for TNFR-Wgn, particularly in physiological conditions, in the transduction of TNF-Egr activity, became controversial. Here we show a role for TNFR-Wgn during normal development, independent of TNF-Egr, in regulating the activity of the FGFR-Btl.

In contrast to TNFR-Grnd which localises to the apical membrane ^17,22^, we find that TNFR-Wgn does not stably localise to the membrane of the tracheal cells and is found instead in intracellular vesicles. This unusual localisation for a receptor seems to be a general feature of TNFR-Wgn, as it was described to be localised in intracellular vesicles also in imaginal tissues ^22^. We observed this same pattern in other tissues in which TNFR-Wgn is expressed (Fig S3D). Our analysis indicates that these intracellular vesicles mostly correspond to endosomes. When endocytosis is generally compromised, TNFR-Wgn is stabilised at the membrane, indicating that in normal conditions the receptor is rapidly internalised after reaching the membrane. In addition, when TNFR-Wgn internalisation is compromised, the capacity of TNFR-Wgn to prevent terminal cell differentiation is lost. Thus, we propose that TNFR-Wgn is constitutively internalised, as it is the case for transferrin receptors ^38^, and that this internalisation is absolutely required for its activity. Future analysis of the intracellular domain of TNFR-Wgn should help to identify the signals that promote this constitutive internalisation. Interestingly, TNFR-Wgn contains a dileucine motif in the intracellular domain, which is not present in TNFR-Grnd receptor and could act as a recognition motif for internalisation ^39,40^.

Our co-immunoprecipitation experiments show that TNFR-Wgn forms a complex with the FGFR-Btl. It was previously shown that TNFR-Wgn can also physically interact with Moesin in the context of photoreceptor axon guidance ^20^. Thus, TNFR-Wgn has the ability to bind diverse proteins, besides its canonical adaptor protein Traf2 ^15^, and thereby regulate their activity. Interestingly, not only TNFR-Wgn, but also the other *Drosophila* TNFR, TNFR-Grnd, was shown to be able to bind an unrelated protein, Veli ^17^. Strikingly, Fn14, a rat TNFR superfamily member, was shown to physically interact with FGFR1 ^41^. Altogether these results indicate that TNFR members have the ability to bind unrelated proteins, and we propose that binding unrelated proteins and regulating their function may be a general mechanism of activity for TNFRs. The participation of TNF-TNFRs in cancer and inflammatory diseases is well-documented, and TNF-therapies directed to control their activity have been developed, but improved therapies are needed to be more effective and avoid undesirable side effects ^6,7^. A better understanding of the molecular mechanisms of TNFRs is key to developing improved therapies.

## Acknowledgements

We thank N. Martín for technical help and J. Ferrandiz for contributions at the initial stages of this work. We thank K. Basler for kindly providing the Wgn antibody, and S. Roy for kindly providing *bnl* and *btl* tagged-alleles. We also thank M. Milan, T. Tanaka, L. Jiang, S. Hayashi and D. Andrew for kindly providing flies and antibodies. We acknowledge the Bloomington Stock Centre and the Developmental Studies Hybridoma Bank for fly lines and antibodies. We thank the members of the Llimargas and Casanova labs for helpful discussions. We thank P.A. Lawrence for help, support and advice, and J. Casanova and M. Furriols for critical reading of the manuscript.

## Author contributions

Conceptualization: AL,ML

Methodology: AL

Investigation: AL,MLE,ML

Formal Analysis: AL,MLE,ML

Supervision: ML

Writing—original draft: ML

Writing—review & editing: AL,MLE,ML

Funding acquisition: ML

## FIGURE LEGENDS

**Fig S1:**
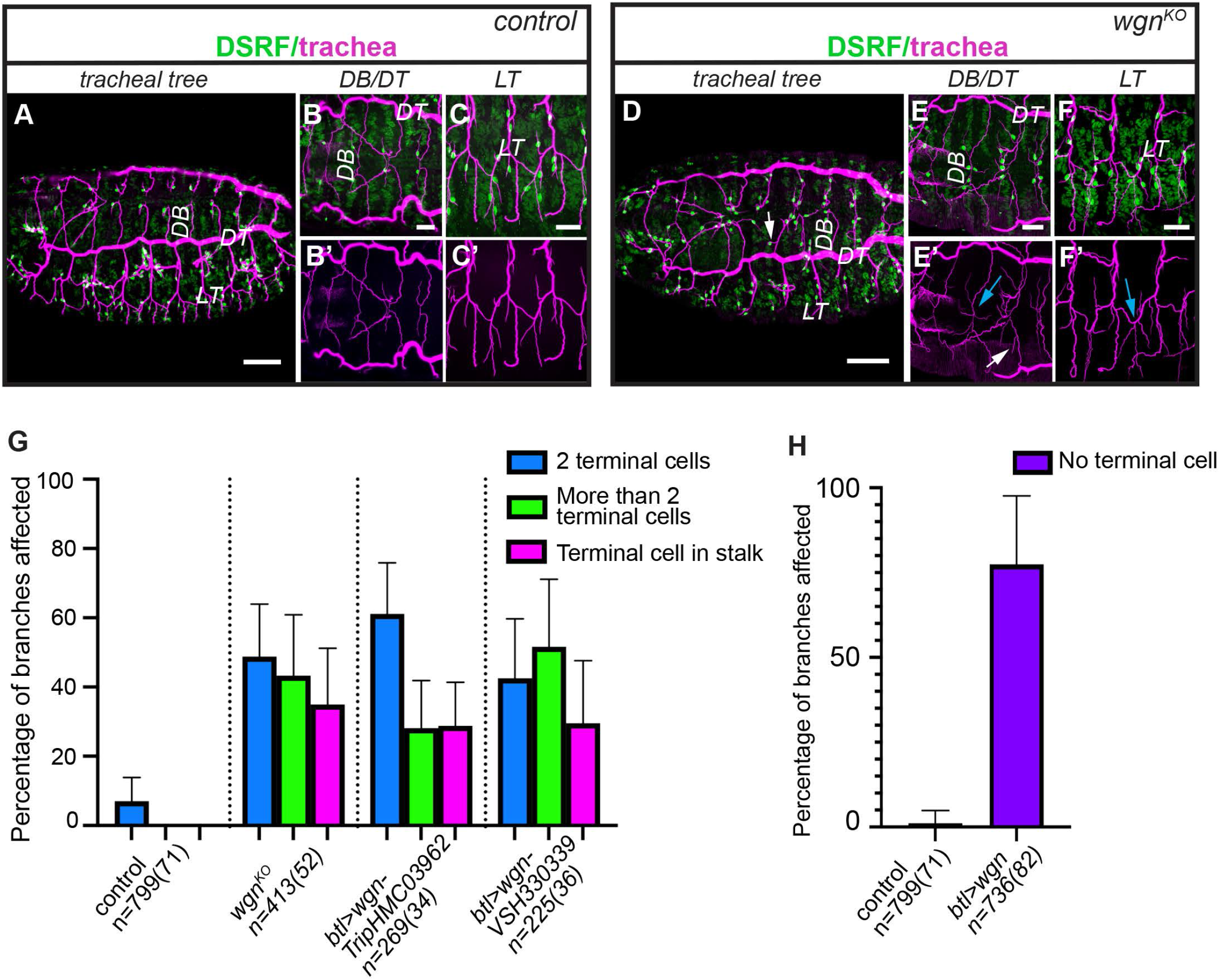
Analysis of the tracheal requirements of *TNFR-wgn*. (A-F) Control or *TNFR-wgn*_ mutant embryos at late stage 15-stage 16 stained with CBP to visualise the tracheal tubes (magenta) and with DSRF to visualise the terminal cells (green). Different views show an excess of terminal cells in *TNFR-wgn* mutants compared to control, which generate excess of terminal branches (white arrows point to terminal branches in the DB stalk). In spite of these defects, the general branching pattern and the fusion events (LT,DB, blue arrows in E’, F’) are not affected (G,H) Quantification of the percentage of dorsal branches that show the indicated phenotypes (excess of terminal cells in *TNFR-wgn* loss of function mutants and lack of terminal cells in *TNFR-wgn*_overexpression) in the different genotypes. Note the high expressivity of the phenotypes. All embryos analysed showed defects (100% penetrance). Error bars indicate standard deviation (s. d.). n, number of DBs analysed, in brackets number of embryos analysed. Scale bar: A,D 50 μm; B,C,E,F 20 μm

**Fig S2:**
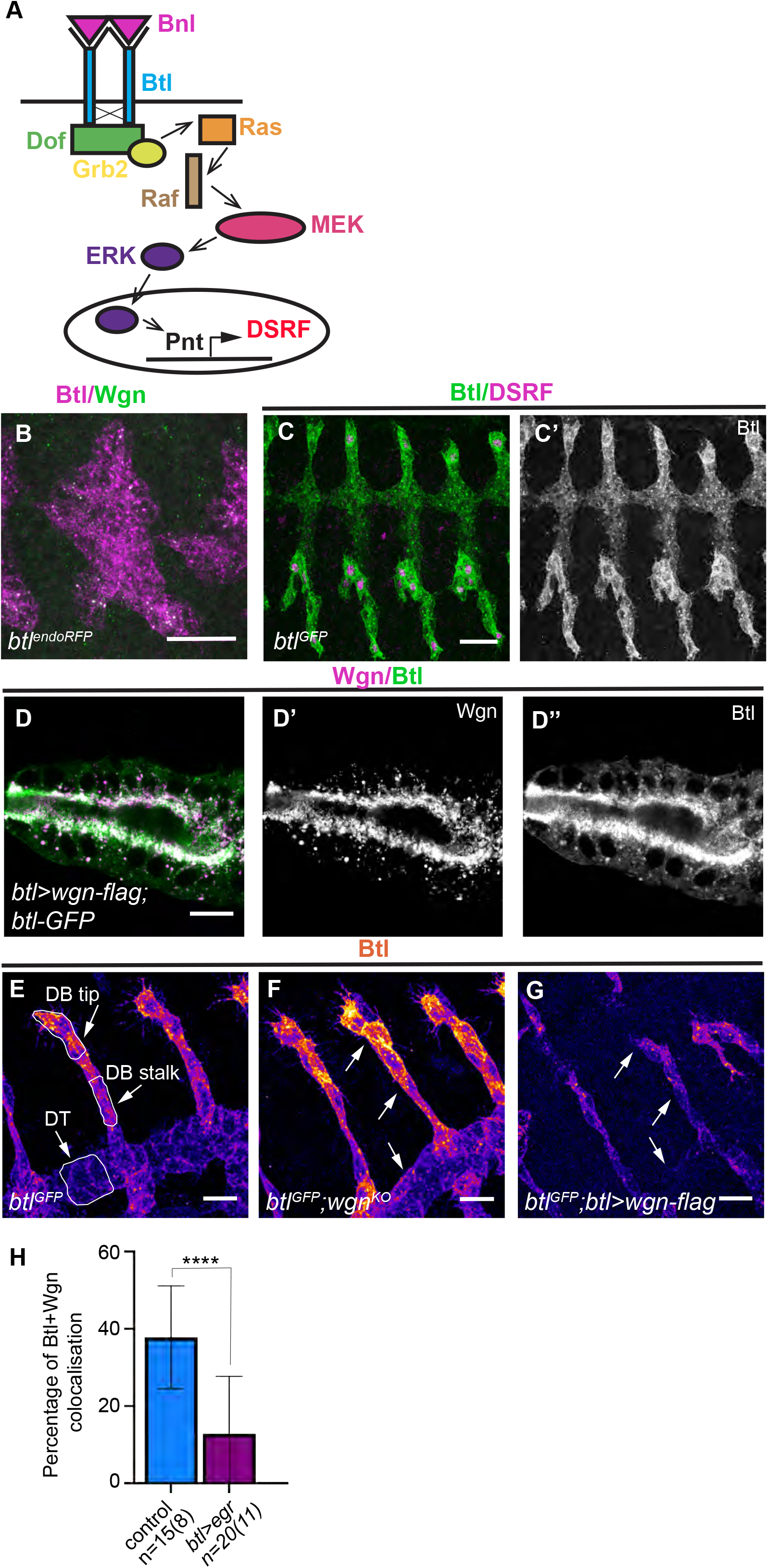
FGFR-Btl pathway in terminal cell specification. (A) Scheme of the FGFR-Btl pathway (B,C) Lateral views of stage 11/12 (B) and 14 (C) embryos showing the pattern of FGFR-Btl accumulation using tagged alleles. Note the increased accumulation of FGFR-Btl at the tips, in DSRF expressing cells, at stage 14 (C’). (D) Salivary gland expressing *TNFR-wgn-Flag* and *FGFR-btl-GFP* showing the accumulation of the two proteins in common intracellular vesicles (E-G) Lateral views of stage 14 embryos showing a representative example of the accumulation of FGFR-Btl in the different genotypes indicated. Regions used for quantifications in Fig 3K are shown. The fluorescence intensity of *FGFR-Btl* is shown in heat maps. (H) Quantification of the percentage of vesicles that contain TNFR-Wgn and FGFR-Btl in the indicated genotypes. Scale bar: B,C 20 μm; D-G 10 μm

**Fig S3:**
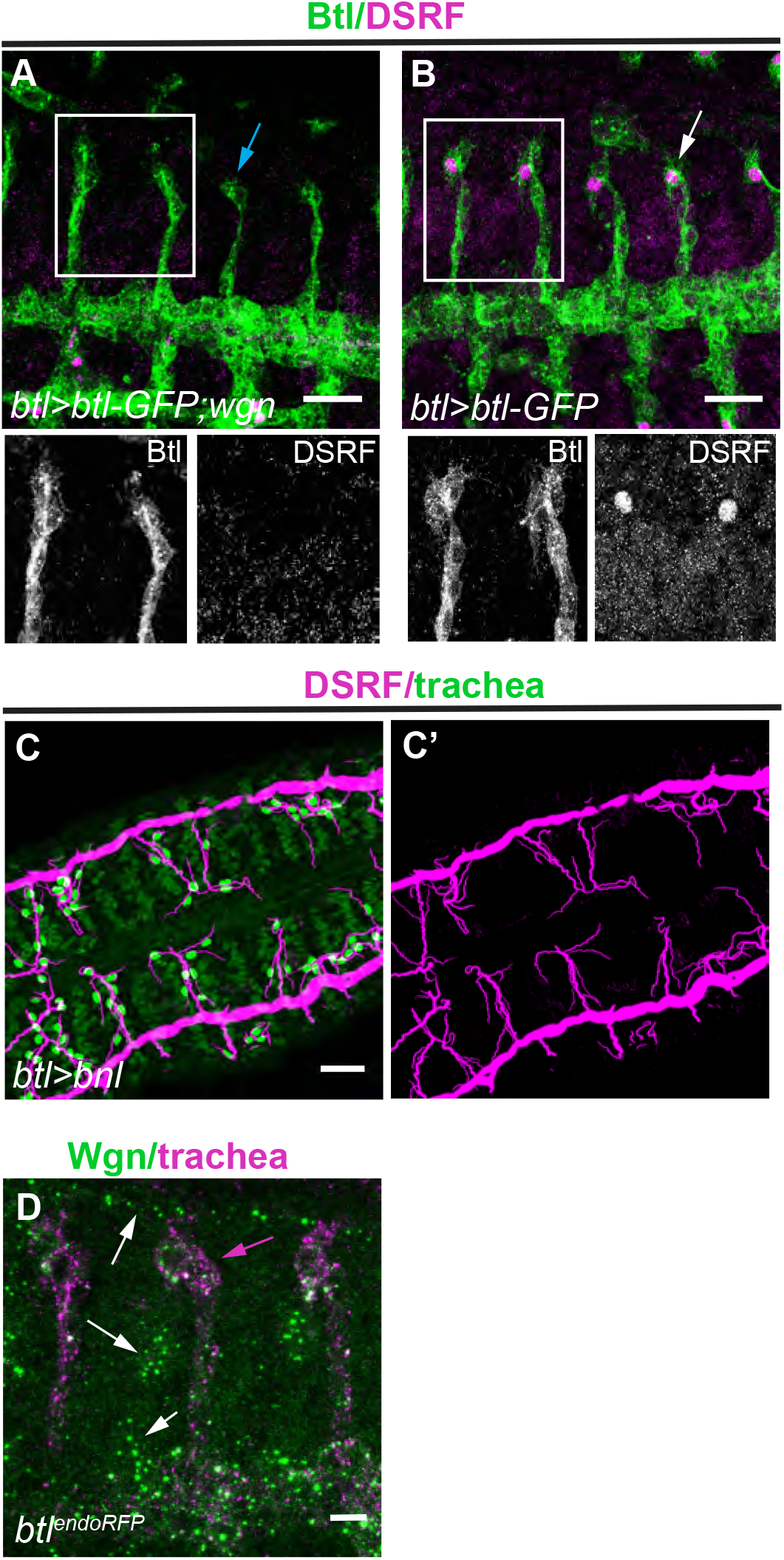
The role of FGFR-Btl pathway in terminal cell specification. (A,B) Lateral views of stage 14/15 embryos of the indicated genotypes stained with GFP (green) to visualise the *FGFR-btl-GFP* overexpression and with DSRF to detect the terminal cells. When *TNFR-wgn* is overexpressed no terminal cells are specified (blue arrow in A) in spite of presence of overexpressed *FGFR-btl*. *FGFR-btl* overexpression does not lead to extraterminal cells (white arrow in B). (C) Lateral view of a stage 14/15 embryo expressing FGF-Bnl in tracheal cells stained with DSRF (magenta) and CBP (green). FGF-Bnl tracheal expression leads to many extra terminal cells. (D) Lateral view of a stage 14 embryo stained with αWgn (green) and with RFP to visualise the tracheal cells (magenta, magenta arrow). Note the presence of TNFR-Wgn in intracellular vesicles in tissues other than the trachea (white arrows) Scale bar: A-D 20 μm

## STAR METHODS

### Key Resources Table

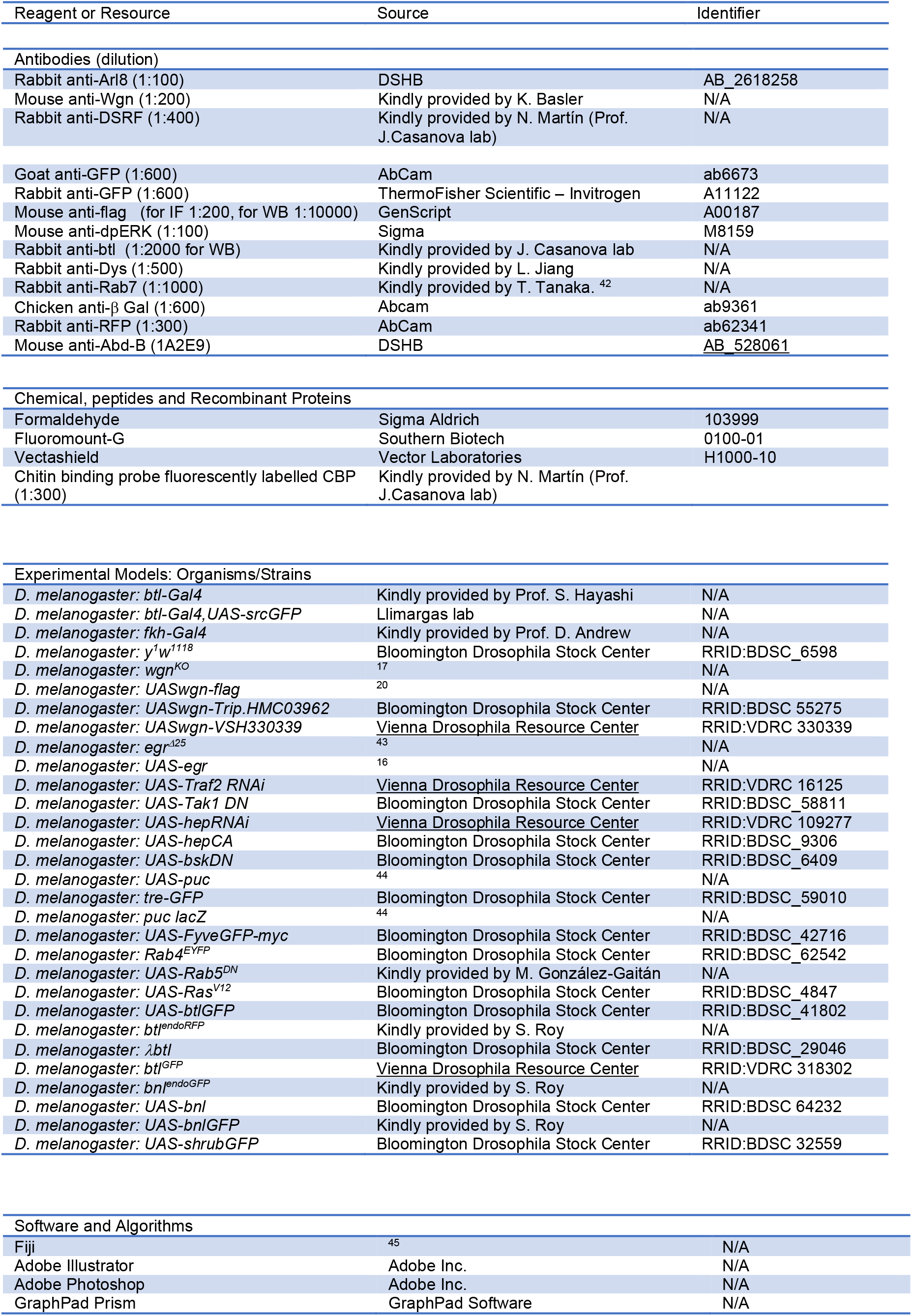

### Resource availability

Detailed information and requests for resources generated in this study should be addressed to and will be fulfilled by the lead contact Marta Llimargas.

All data reported in this paper and any additional information required to reanalyze the data described in this paper is available from the lead contact upon request.

This paper does not report original code.

### Experimental model and subject details

#### *Drosophila* strains and maintenance

All *Drosophila* strains were raised at 25°C under standard conditions. Balancer chromosomes were used to follow the mutations and constructs of interest in the different chromosomes. For overexpression experiments, we used the Gal4 drivers *btlGal4* (in all tracheal cells) and *fkhGal4* (in salivary glands). The overexpression experiments were performed using the Gal4/UAS system ^46^. To maximise the expression of the transgenes, crosses were kept at 29°C. The fly strains used are listed in the “Resource table: Experimental models”.

### Methods details

#### Immunohistochemistry

Embryos were stained following standard protocols. Embryos were staged as described ^47^. Embryos were fixed in 4% formaldehyde (Sigma-Aldrich) in PBS1x-Heptane (1:1) for 20 min. Embryos transferred to new tubes were washed in PBT-BSA blocking solution and shaken in a rotator device at room temperature. Embryos were incubated with the primary antibodies in PBT-BSA overnight at 4°C. Secondary antibodies diluted in PBT-BSA (and for the CBP staining) were added after washing and were incubated at room temperature for 2–5 h in the dark. Embryos were washed, mounted on microscope glass slides and covered with thin glass slides. The primary antibodies used are listed in the “Resource table: Antibodies”. Cy3-, Cy2- and Cy5-conjugated secondary antibodies (Jackson ImmunoResearch) were used at 1:300.

#### Image acquisition

Images from fixed embryos were taken using Leica TCS-SPE or Leica DMI6000 TCS-SP5 laser confocal microscopes, with the 20x and 63x immersion oil (1.40-0.60; Immersol 518F – Zeiss oil) objectives and additional zoom. Settings were adjusted for the different channels prior to image acquisition. Z-stack sections of 0.24-0.5 μm were acquired. The images were imported and processed using Fiji (ImageJ 1.49b) for measurements and adjustments, and assembled into figures using Photoshop and Illustrator.

#### Image analyses

##### 1. Quantification of terminal cells

The number of terminal cells in dorsal or ganglionic branches was calculated using the nuclear factor DSRF as a marker for the nuclei of terminal cells. CBP was used to mark the lumen of the tracheal system to identify the different branches. The Max Intensity projections of confocal sections of late stage 14-stage 15 embryos, from different immunostaining experiments, were analysed using Fiji. DSRF positive nuclei were manually selected with the wand tool in Fiji and counted for each branch/embryo.

##### 2. Quantification of vesicles

To quantify the number of vesicles, Max Intensity projections of late stage 14 embryos were taken and analysed using Fiji. After substracting the background, a Region Of Interest (ROI) was drawn to select the dorsal branch. A binary mask was created using the threshold tool and the watershed segmentation tool. Number of vesicles were counted using the Analyse particles tool and the parameters were set to 0,05-1,7 mm^2^ size, 0-1 circularity; the number of vesicles and a mask of the result were obtained.

##### 3. Quantification of levels

To analyse the levels of Btl protein in the tracheal cells and to compare control and *wgn* mutant conditions we performed different independent experiments in which control and mutant embryos were collected, fixed and stained together. Confocal images of late stage 14 embryos were acquired with the same laser settings for each individual experiment. We then generated a projection from the different stacks using the Max Intensity tool in the Fiji software and subtracted background. Three different ROIs were considered and compared: the tip of a dorsal branch, a part of the stalk of the same branch and a part of the dorsal trunk near the dorsal branch. To measure the total Btl fluorescence at each ROI we obtained the “integrated density” in manually drawn areas at the tip, stalk and adjacent dorsal trunk with the freehand selection tool (see Fig S3E). The integrated density of each region was normalised to the average of the integrated densities calculated for the corresponding region (tip, stalk and dorsal trunk) in the control of each experiment. The obtained values were compared between control and *wgn* mutant conditions using the Scatter Plot tool of GraphPad Prism.

##### 4. Colocalisation of vesicles

Colocalisation analysis was performed using the ImageJ plugin Colocalisation highlighter, considering colocalisation when the ratio of fluorescence intensities between the two channels analysed was above 0,5. Those fluorescence intensities above the threshold appear in a binary image colour as white (colocalised points). From this mask, we selected manually each vesicle with colocalisation with the wand tool in Fiji and added it in the ROI Manager to be counted.

### Co-immunoprecipitation assay

Assays were performed with extracts prepared from salivary glands of *Drosophila* third-instar larvae that were lysed in RIPA buffer (50 mM Tris-HCl pH8,150 mM NaCl, 0.1% SDS, 0.5% sodium deoxycholate,1% Triton X-100, 1mM PMSF and protease inhibitors (cOmplete Tablets, Roche). Extracts were immunoprecipitated using αFlag antibodies or a control antibody (αAbd-B), followed by incubation with Protein G Dynabeads (Invitrogen). Immunoprecipitates were washed with RIPA buffer and analysed by Western blot using either αBtl or αFlag antibodies and the Immobilon ECL reagent (Millipore).

### Quantification and statistical analysis

Data from quantifications was imported and treated in the Excel software and/or in GraphPad Prism 9.0.0, where graphics were finally generated. Graphics shown here are scatter dot plots or columns, where bars indicate the mean and the standard deviation. Statistical analyses comparing the mean of two groups of quantitative continuous data were performed in GraphPad Prism 9.0.0 using unpaired two-tailed student’s *t-test* applying Welch’s correction. Differences were considered significant when p < 0.05. Significant differences are shown in the graphics as: *p < 0.05, **p < 0.01, *** p < 0.001, **** p < 0.0001. Sample size (n) is provided in the figures or legends.

